# Nutrient learning, perception and generalization differ among wild pollinator species

**DOI:** 10.1101/2023.08.04.551969

**Authors:** Ana Laura Pietrantuono, Valeria Fernández-Arhex, Jean-Christophe Sandoz, Fabrice Requier

## Abstract

Learning is the process through which skills, knowledge, behaviors are acquired and developed. Through life experiences, pollinator insects, learn to associate odor or visual stimuli from flowers with a rewarding food as pollen or nectar. This capability allows them to obtain resources efficiently and provides a valuable pollination service. Here, we compared the learning capability among different wild pollinator insects by mean of experiments based on the recent Free-Moving Proboscis Extension Response (FMPER) technique. Specifically, we evaluated the process of perception, learning and generalization between pollen enriched with different concentrations of fatty acid (which play a critical role in the survival, development and reproduction of many animals). We compared the associative learning in two castes of bumble bees (queens and workers of *Bombus terrestris*), honey bees (*Apis mellifera)*, solitary bees (*Ruizanthedella mutabilis*) and non-bee pollinator species (the hoverfly *Eristalis tenax*). Our results reveal that learning performances differed among bumble bee social castes, with queens acquiring learning odorants more effectively than workers. Likewise, learning performances differed among the four species of insect pollinators. Honey bees acquired odor-sucrose association more rapidly than the other species. Likewise, bumble bees learned better than the solitary bee species and the hoverfly. The pattern of generalization among odorant stimuli as also different among the studied species, with honey bees showing stronger generalization and hoverflies showing more specific response patterns. Studying the learning behavior in insect pollinators provides valuable information for the conservation of these species and services they provide, through adapted pollinator-friendly schemes matching their behavioral performances.

## Introduction

Insects play a critical role in agro-ecosystems providing significant services for nature and human well-being [1,2]. Bees, butterflies, wasps, flies and beetles represent the most important insect pollinators in croplands and (semi-)natural ecosystems [3–6]. However, populations of insect pollinators are declining in many parts of the world mainly due to agricultural intensification [7–9]. In particular, land use change and the increased use of inputs (e.g. herbicides, fertilizers) have altered the availability and quality of floral resources [10–12] on which pollinator fitness is dependent [13].

Flowers offer the two main food resources for insect pollinators. Nectar mostly provides carbohydrates, while pollen provides proteins and fatty acids, which constitute the main food supply for offspring development [14–17]. During their foraging lifetime, insect pollinators face to a wide diversity of plants that vary in their quality, nutrient profile [18–20] and also in the release mixtures of volatile substances that produce. Insects learn to associate these complex and diverse odor cues with food rewards. This capability to learn, perceive and generalize among the diversity in floral resource qualities is assumed to directly affect the ability of insects to survive in an ecosystem, with critical implications for conservation [20–23]. Previous studies demonstrated that insect pollinator species could have different cognitive process, mainly related to their pattern of foraging behavior, degree of socialization and even brain size [24–28]. One of the most studied insects from a behavioral perspective, with a history of classical conditioning experiments and greatest cognitive capability is the Western honey bee [29–31]. Nevertheless, very little is known about the associative learning behavior among other pollinator species. Learning comparisons among different wild pollinator insects is fundamental for a better understanding of complex insect-plant interactions and above all of their behavior and the ecological role they play. Here, we assessed the capability of different wild-living insect pollinator species to learn, different pollen resource qualities, since we considered that the way in which the odorant of pollen are perceived could affects the behavior and success of an insect. Particularly, we focused in fatty acids which play a critical role in the survival, development and reproduction. Fatty acid act as a compact form of energy storage and as components of cellular and subcellular bio-membranes [32]. In insects, these nutrients can be used to fuel flight and contribute to the biosynthesis of pheromones, eicosanoids, waxes, and as components of defensive secretions [33–35]. Consumption of fatty acids can be fundamental for insects’ success, a low intake can reduce learning performances in honey bees [36] while a high intake can reduce reproduction and survival [37].

In the present study, we examined the learning behavior of four species of wild pollinators insects. Specifically, we asked whether this capability differ among species and if it depends on the concentration of fatty acids on the stimulus. To answer these questions, we performed PER conditioning and evaluated generalization among the different species and pollen stimuli (pure pollen and pollen enriched with different concentrations of stearic acid). We carried out field realistic experiments in North Patagonia (Argentina) on two social species, *Apis mellifera* and *Bombus terrestris* (Hymenoptera: Apidae), and on two solitary species for the first time. Specifically, the solitary bee *Ruizanthedella mutabilis* (Hymenoptera: Halictidae) native to Argentina, and the hoverfly *Eristalis tenax* (Diptera: Syrphidae) exotic in Argentina (native to Europe). Both solitary species are known as important pollinators [38–39] but have been little studied so far in terms of their learning capabilities and behavior. In addition to this inter-specific aspect, we also assess the learning process of two social castes in *B. terrestris,* i.e. queen and worker. For all species and castes, we performed conditioning experiments using the recently developed technique of Free-Moving Proboscis Extension Response (FMPER; [40]).

We expected different degrees of learning performances among the studied pollinator species. First, it was recently shown that body size affects learning capability (i.e. larger body sizes suggest better learning capabilities; [26]). Thus, considering that the Western honey bee and *B. terrestris* are the species with the larger body sizes among the insects studied here, and that they also possess big brains [41], we argued that they will learn more quickly than the solitary bee. Among bumble bee queens and workers, we assumed that queens would show better learning performances than workers for the same reason (queens have larger body size than workers) but also because queens play a critical role for colony development and their learning abilities must support quick and efficient food collection [42]. Finally, we considered that the life-history trait, as the level of sociality which influence learning abilities (i.e. sociality per se favors the evolution of learning - The social brain hypothesis-[24]) could also determine learning performances among pollinators species. Thus, we expected that the level of learning would be higher for social bees, followed by solitary bees and finally non-bees pollinators.

## Materials and Methods

The study was carried out in the proximity of the city of San Carlos de Bariloche (41°04′12″S 71°09′54″W, 893 m a.s.l.), Río Negro, Argentina. This region is an ecotone zone between the Andean-Patagonian forests and the steppe, with a temperate-cold climate and a West–East rainfall gradient generated by the Andes Mountains. It has a wide diversity of floral resources composed of native and exotic herbs, shrubs and trees. Data collection and experiments were started at the end of summer (March - April) 2020 and continued during the spring and the next summer season (September 2020 - April 2021).

### Insect pollinator species

We collected four species of insect pollinators: the Western honey bee *Apis mellifera*, the bumble bee *Bombus terrestris*, the solitary bee *Ruizanthedella mutabilis* and the hoverfly *Eristalis tenax*. All of these insect pollinators were collected from wild flowers between 11 a.m. and 4 p.m. using a handheld insect aspirator (see [43]). Only female individuals were collected. We differentiated queens and workers of *B. terrestris* based on body size. Queens are markedly larger and more than three times heavier than the largest workers [44–45]. In total, we collected 102 (worker) individuals of *A. mellifera*, 134 individuals of *B. terrestris* (n=60 queens, n=74 workers), 72 individuals of *E. tenax* and 88 individuals of *R. mutabilis*. Once captured, each individual was placed in a plastic tube with a cap (15 cm long × 2.5 cm diameter) for the FMPER experimental procedure. The tubes had two perforations (2 mm diameter) at the curved end; it is through these perforations that the stimuli were introduced during the learning procedure (see below-Learning protocol). The experiment did not require any anesthetization or immobilization of the insects. Prior to the beginning of the learning procedure (approximately 10 min before), we evaluated each insect’s appetitive motivation by offering a toothpick soaked in sucrose solution (50% v/v) into the tube. Individuals that did not perform an extension of the proboscis were discarded from the experiment (see Data analysis). Following the procedure in [42], individuals were then allowed to acclimatize to the tube for 1 h (bees and hoverfly) and 2 h (*B. terrestris*) in total darkness at a temperature of 20 ± 2°C. We stablished these resting periods based on previous studies [42,46–47] and are assumed to reduce stress from the capture and also to increase appetitive motivation through a period of starvation. At the end of this phase, the insects were found to be calm and motivated to participate in the experiment.

### Pollen enriched with Fatty Acids (FA)

Samples of pollen pellet were collected from honey bee hives at the field station of the National Institute of Agricultural Technology – Bariloche Research Lab (EEA INTA Bariloche, San Carlos de Bariloche, Province of Río Negro, Patagonia, Argentina, 41° 7’ 22.673’’ S, 71°15′ 04.94″ W), located within the study area. Honey bee pollen collection was performed using pollen traps (Apipolen®, Madrid, Spain) on three hives during 24h on three austral summer dates (10 February, 15 February and 2 March of 2017). All the samples were left to dry at room temperature (22°C) over 48 h and then pooled. The pooled sample was divided in four sub-samples of 8 grams, with one for determining the botanical origin and the rest for stimuli preparation (see below). With one of these sub-samples we proceeded to classify the pollen pellet by color and then we selected the five most abundant (which represented 76% of the total weight of the sample). We then took one pellet of pollen of each color and observed under a scanning electron microscope (SEM) the morphological characteristics of the pollen grains [23, 48]. We also took high-resolution photographs of individual pollen grain structure and this information was compared with previous samples and bibliographical references allowing the determination of the botanical origin of the loads [49–51]. Photographs were captured at the Atomic Center of Bariloche (Centro Atómico Bariloche, Depto. Caracterización de Materiales – Servicio de Microscopia y Rayos X, San Carlos de Bariloche, Río Negro, Argentina).

In addition, post-identification validation was carried out by verifying the presence of the identified plant species in the landscape surrounding the hives. We found three dominant plant species which belong to the Brassicaceae family, one of the main sources of pollen for honey bees in the study region. Brassicaceae is a stenopalynous family (i.e. the palynological characteristics analyzed do not allow differentiation among the studied species) and it was not possible to differentiate its genera based on the observation of the pollen loads. We determined three morpho-types: Type *Diplotaxis* sp. (possibly *Diplotaxis tenuifolia* and/or *D. virgata*) (30.6%), Type *Sinapis arvensis* (25.7%) and Type *Raphanus raphanistrum*, possibly *Arabidopsis thaliana* or *Lepidium draba* (19.7%). The rest of the sample was made up of *Foeniculum vulgare* (10.5%) (Apiales: Apiaceae) and *Sambucus nigra* (9.1%) (Dipsacales: Adoxaceae).

The three remaining sub-samples were used to produce the three conditioned stimuli for the learning experiment. One sub-sample corresponded to the control or pure pollen, i.e. it was not enriched with stearic acid. It is hence called 0×FA (for Fatty Acid). The two other sub-samples were enriched with stearic acid, producing one solution of slightly enriched pollen (0.5×FA) and another of highly enriched pollen (10×FA). For the stimuli preparation we grounded the sub-samples in a mortar until we obtained a very fine and homogeneous powder. Then we added 5 mL of distilled water to each one and the corresponding amount of stearic acid for the enriched solutions. Amounts of stearic acid were determined based on previous studies [37, 52] which analyzed the fatty acid content among various plant species and calculated that the natural concentration of stearic acid is on average 2.46 mg/g. Thus, based on these references we added (per gram of pollen powder) 1.23 mg of stearic acid for the slightly enriched pollen (0.5×FA) and 24.6 mg for highly enriched pollen (10×FA). We finally obtained a paste with a creamy consistency that sticks to the toothpick that we would use to approach the stimuli to the insects (see S1.1 Figure in Supporting information for more details).

### Free-Moving Proboscis Extension Response (FMPER)

To study the learning abilities of the four insect pollinators, we used the recently developed “Free-Moving Proboscis Extension Response (FMPER)” protocol [40]. The FMPER technique is based on the natural and innate Proboscis Extension Response (PER) performed by several species of insects like bees when they land on flowers and detect the presence of nectar with their antennae [29, 53]. In classical conditioning of the proboscis extension in honey bees the procedure is performed under laboratory conditions, mostly using harnessed honey bees that have been previously anesthetized [31]. A challenge for performing FMPER is that the experimenter has less control over the learning variables (time intervals and durations of CS and US). On the other side, FMPER offers the great advantages to not anesthetize the insects, to let the individual free to move in the tube, and to be applied in the field with wild species without much manipulation. Overall, individuals are less stressed and can be released again after conditioning. This represents an important point when considering native wild pollinators in a context of pollinator decline [7–9].

### Learning protocol

#### Conditioning phase

After the acclimatization period, a conditioning phase was applied (adapted from [23]). Insects were trained to associate an olfactory stimulus, i.e. a pollen solution (Conditioned Stimulus - CS) with a reward of sucrose solution - 50% v/v (Unconditioned Stimulus - US). The procedure consisted of three conditioning trials (C1, C2 and C3), each trial lasting one minute with an inter-trial interval (ITI) of 10 minutes. During each trial we first introduced the CS through one of the perforations at the end of the tube, and when the insect had its body oriented to the stimulus and began to walk in that direction, we started the clock to measure the duration of the trial (1 min). When the individual was close enough to the toothpick and oriented both its antennae (in the case of bees) or moved its front legs (in the case of Diptera) we approach the CS to the anterior part of the insect’s body, allowing the individual to perceive odorant stimuli. This toothpick was immersed with the CS (i.e. one of the three concentrations of fatty acid: 0×FA; 0.5×FA; 10×FA). After 15 seconds, the US was introduced into the tube and presented to the anterior part of the insect’s body for 10 seconds. If during this period of time a PER was observed, we brought the US toothpick to the insect’s mouthparts allowing it to consume the sucrose). After that, both toothpicks were removed simultaneously from the tube (see S1.2 Figure for Learning protocol). At each trial, the introduction (left or right perforation on the tube) of the rewarded stimulus was alternated and the toothpick with the stimulus was renewed.

We registered the insect behavior after the presentation of each stimulus (CS and US) and considered a positive response to the stimulus when the individual performed a full extension of the proboscis (PER, which includes a wide opening of the mandibles and protrusion of the proboscis) during at least 5 seconds in the direction of the stimulus. The behavior of Diptera was challenging to record mainly due to the small size of the insect. In this case, we evaluated the orientation of the first legs approaching the stimuli. It should be noted that during the learning protocol, the tubes were attached to a plastic container in order to minimize movements of the tube and unnecessary stress for the insect (see S1.3 Figure for diagram device).

#### Test phase and generalization

The test phase began 10 minutes after the conditioning phase and consisted in a sequence of three trials with ITIs of 10 minutes. At each trial, one of the three pollen stimuli was presented but without reward, in a random order, during 10 seconds. The goal of this phase was to evaluate whether insects would respond to the learned pollen stimulus and to novel stimuli (i.e. a type of pollen different from the one used for conditioning). If an individual extends its proboscis when presented with a novel stimulus, it indicates that it is perceived as similar to the CS, thus providing evidence of generalization.

Finally, we re-evaluated the motivation of each individual 10 min after the end of the experiment to determine whether their motivation was maintained by offering again a toothpick soaked in sucrose solution into the tube. All of the individuals responded to the sucrose and no individual was discarded for this reason from the data analysis (see below). The assays were always carried out in series of 10 individuals and at the end of the protocol the tubes were first washed with detergent (odorless, colorless and with a neutral pH) and then rinsed with plenty of water and finally with ethyl alcohol and allowed to dry. To prevent recapture of insects, each individual was marked prior to release with a dot of washable paint on the thorax [54].

### Data analysis

Individuals were randomly assigned to one of the different treatments (one treatment per pollen solution) and each individual was used only once in the experiment. Among the total number of individuals used (n=396), some were discarded from data analysis, following standard criteria [55]: 1) if it responded with a PER to the CS at the first training trial, as it could mean that the insect had already formed a pollen odor-sugar association; 2) if it never responded with PER to the US during all of the conditioning trials, as this suggests a lack of appetitive motivation, precluding conditioning. Overall, 53 individuals were discarded including one bumble bee queen, one bumble bee worker, 35 honey bee workers, four hoverflies and 12 solitary bees.

All statistical analyses were performed using the R Project for Statistical Computing version 3.6.1 [56]. Binomial general linear mixed models (GLMMs) were used to compare the proportion of PER (response variable) between studied insects, pollen quality categories (fixed factors), the interaction between study insects and pollen quality categories, and trial levels. We also considered the insect identity as random factor in the GLMMs. We used the Akaike Information Criterion (AIC) for selecting the best model (the most parsimonious model in the compromise between fit and complexity) between GLMMs with or without the interaction term of studied insects and pollen quality categories, using the criteria of ΔAIC>2. We selected the less complex model whenever ΔAIC<2 (i.e. the GLMMs without the interaction term). Differences between each pollen quality category and insect species were evaluated with *a posteriori* multiple pair wise comparisons (Tukey’s HSD test). Following the same approach, we compared the proportion of PER between studied insects and pollen quality categories in the Test phase, in order to study generalization. Model residuals were extracted and inspected against fitted values (residuals versus fitted plot and normal Q–Q plot) to ensure residual normality and homoscedasticity assumptions were fulfilled. The significance level for the statistical tests was set at 5% for the risk of rejecting the hypotheses.

## Results

### Conditioning phase

During the course of the conditioning trials the level of PER increased for each insect species and for all pollen treatments. In the case of bumble bees, we could determine that there was no interaction between the studied insects (queens vs. workers) and the pollen quality categories based on the AIC comparison (model with interaction AIC=299.3; model without interaction AIC=295.3). We found that at the end of the conditioning phase (Conditioning trial 3) queen bumble bees showed a higher level of PER for all stimuli than workers (*Z*=2.34, *P*<0.05, **Figure 1A, B**) and that there was a significant effect of the pollen quality categories on PER proportion at the end of the conditioning phase for both castes (Z=4.38, *P*<0.05). The results of the GLMM model performed to evaluate the PER in bumble bees during the Conditioning phase are detailed in S2 Appendix, S2.1 Table (Supporting Information).

When analyzing learning behavior among the four insect species, again the most parsimonious model (ΔAIC>2) was that without any interaction between the studied insects and the pollen quality categories (model with interaction AIC=780.0; model without interaction AIC=777.5). Comparing the proportion of PER between the four insect pollinator species (**Figure 2**), we found that honey bees presented the highest learning success (up to 95% for 10×FA, **Figure 2A**) among all pollinator insects and significantly differed from the other species (**Table 1**). Specifically, significant differences appeared between the honey bee and the solitary bee (*Z*= −5.987, *P*<0.001, **Figure 2B**), the bumble bee (*Z*= −3.310, *P*<0.001, **Figure 2C**) and also with the hoverfly (*Z*= −5.001, *P*<0.001, **Figure 2D**). We also found significant differences between the, bumble bee and the hoverfly (*Z*= −2.647, *P*<0.001) and in the comparison between the bumble bee and the solitary bee (*Z*= −3.987, *P*<0.01). In both comparison, bumble bees reach the higher value of PER for almost all treatments, with the exception that in the third trial of conditioning with 0.5×FA hoverflies have a similar percentage of PER (approx. 82%).

**Figure 1.**
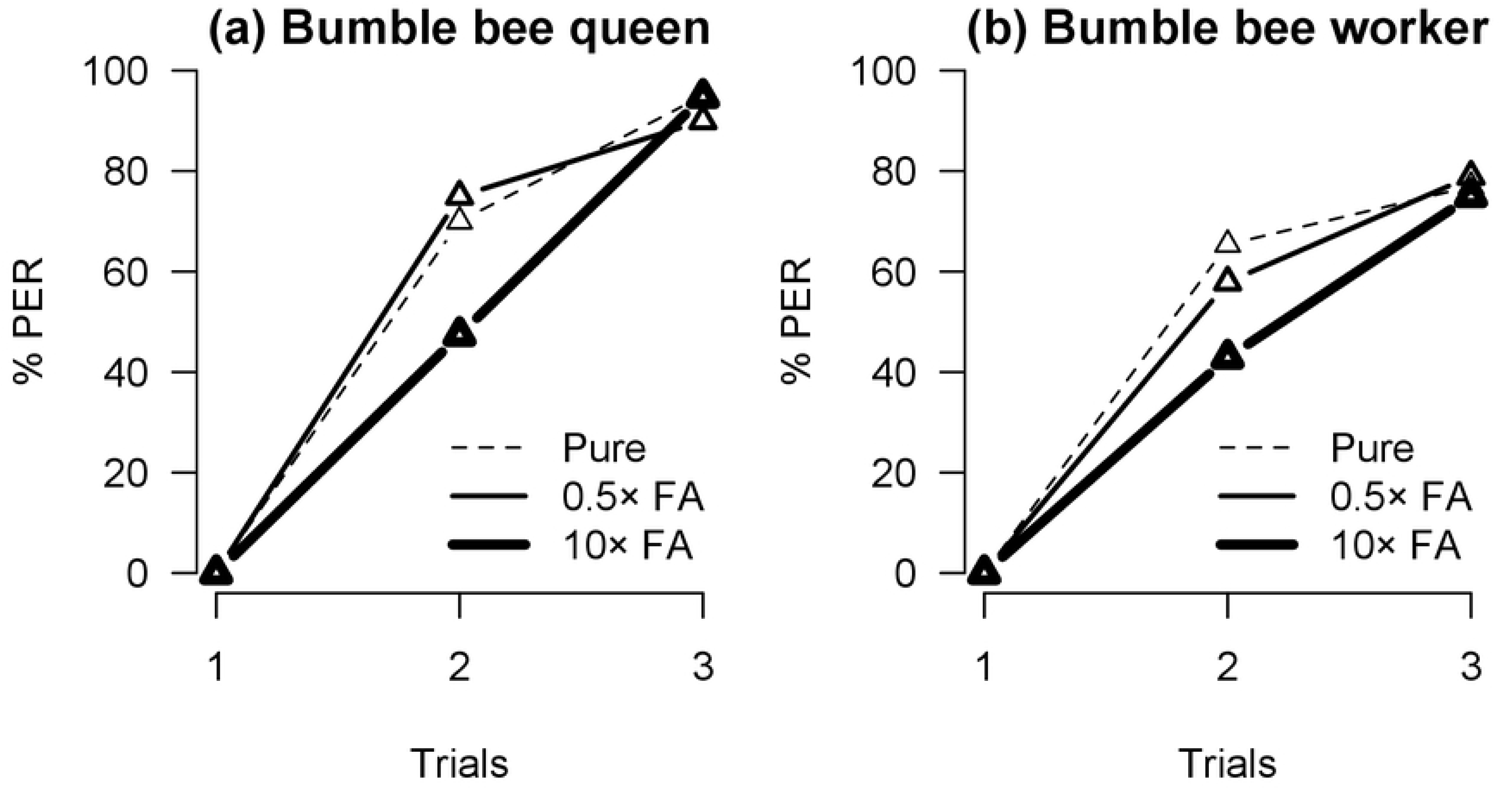
Comparison among (a) bumble bee queens (n=59), and (b) bumble bee workers (n=73) on the percentage of proboscis extension responses (PER) during the Conditioning phase for each pollen quality category (Pure pollen with 0×FA, 0.5×FA or 10×FA) at each conditioning trial (C1, C2, and C3).

**Figure 2.**
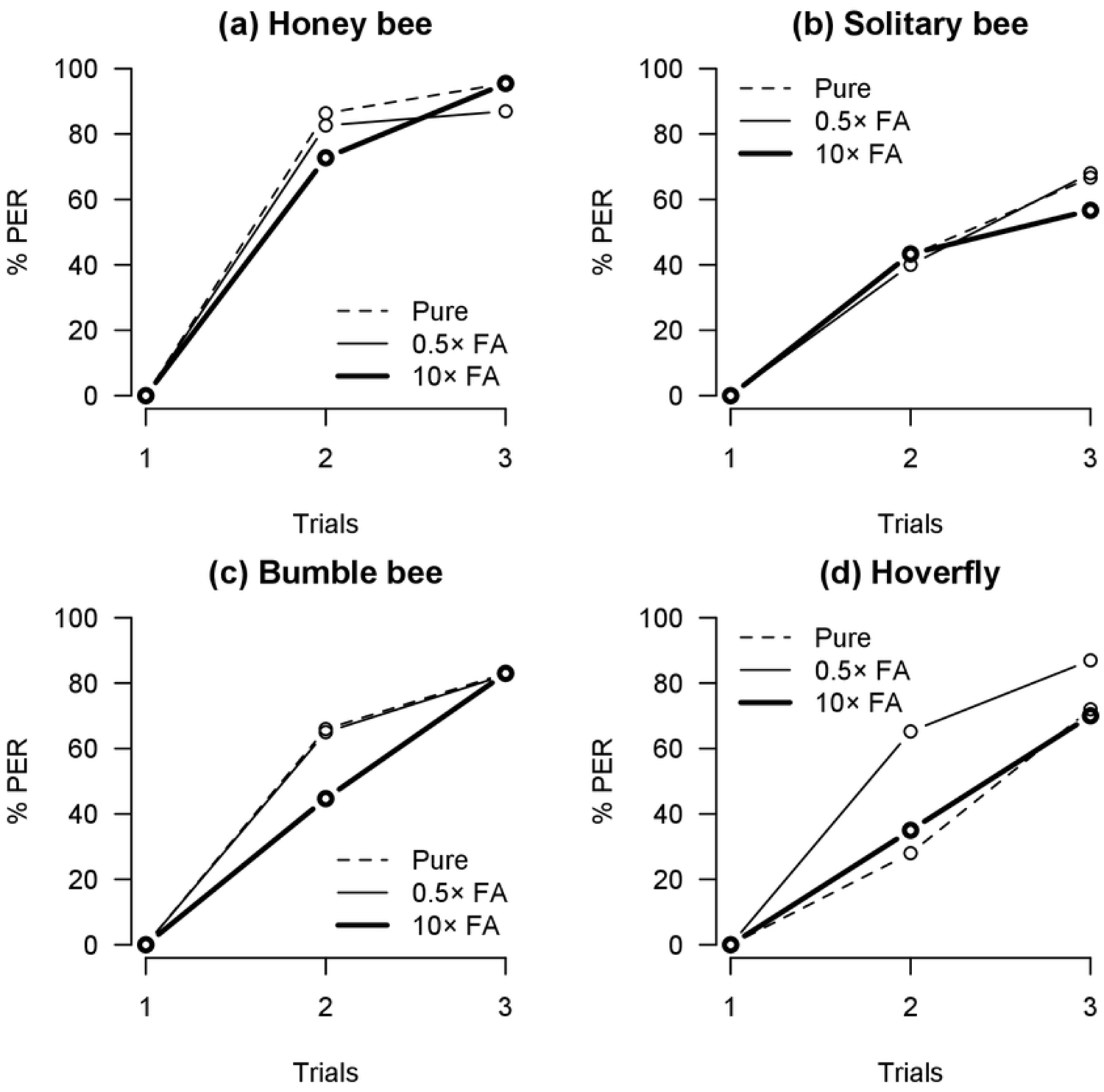
Comparison among (a) honey bee (n=67), (b) solitary bee (n=76), (c) bumble bee (n=132 including both queens and workers) and (d) hoverfly (n=68) on the percentage of proboscis extension responses (PER) during the Conditioning phase for each pollen quality category (Pure pollen with 0×FA, 0.5×FA or 10×FA) at each conditioning trial (C1, C2, and C3).

**Table 1.** Tukey test values comparing the percentage of proboscis extension responses (PER) for different studied insects (Honey bee vs. Bumble bee vs. Hoverfly vs. Solitary bee) from the GLMM testing the effects of studied insects, trial levels and pollen quality categories on PER during Conditioning phase. Bold lines indicate significant differences (P<0.05).

| Parameter | Estimate | Std. Error | z value | *P* value |
| --- | --- | --- | --- | --- |
| **Bumble bee vs. Honey bee** | **-0.9784** | **0.2956** | **-3.310** | **0.00496** |
| **Hoverfly vs. Honey bee** | **-1.5941** | **0.3187** | **-5.001** | **< 0.001** |
| **Solitary bee vs. Honey bee** | **-1.8691** | **0.3122** | **-5.987** | **< 0.001** |
| **Hoverfly vs. Bumble bee** | **-0.6157** | **0.2326** | **-2.647** | **0.03968** |
| **Solitary bee vs. Bumble bee** | **-0.8907** | **0.2234** | **-3.987** | **< 0.001** |
| Solitary bee vs. Hoverfly | -0.2750 | 0.2512 | -1.095 | 0.68822 |

When we evaluated the effects of trial levels (C2 vs. C3) and pollen quality categories on the (PER) during the Conditioning phase for each species, we found significant differences in bumble bee (*Z*= −3.310, *P*<0.001), hoverfly (*Z*= −5.001, *P*<0.001) and solitary bee (*Z*= −5.987, *P*<0.001) (see S2.2 Table). Thus, performances still increased from C2 to C3 in these species, while in honey bees very high performances were already reached at C2, this could indicate that honey bees learn faster compared to other species.

### Test phase and generalization

Figure 3 and 4 present the results obtained during the Test phase between bumble bee castes and between the four species of insects respectively. The AIC comparison of the GLMMs with and without the interaction term between bumble bee castes (queens and workers) and the pollen quality categories resulted in the selection of the model without the interaction term as the most parsimonious models for all the stimuli conditioned. When bumble bees queens were conditioned with 0×FA or 0.5×FA, they responded similarly to all stimuli (**Figure 3A**, S2.3 Table) indicating that were are able to generalize among them. In contrast, when bumble bee queens were conditioned with 10×FA, they responded significant less to the other two stimuli (0×FA and 0.5×FA, *Z*=2.51, *P*<0.05), suggesting that they did not generalize to lower concentrations. On the other hand, when bumble bee workers were conditioned with pure pollen (0×FA), they generalized to all the stimuli. When conditioned with 0.5×FA or 10×FA, the workers generalized only to the other stimulus containing FA (**Figure 3B**).

**Figure 3.**
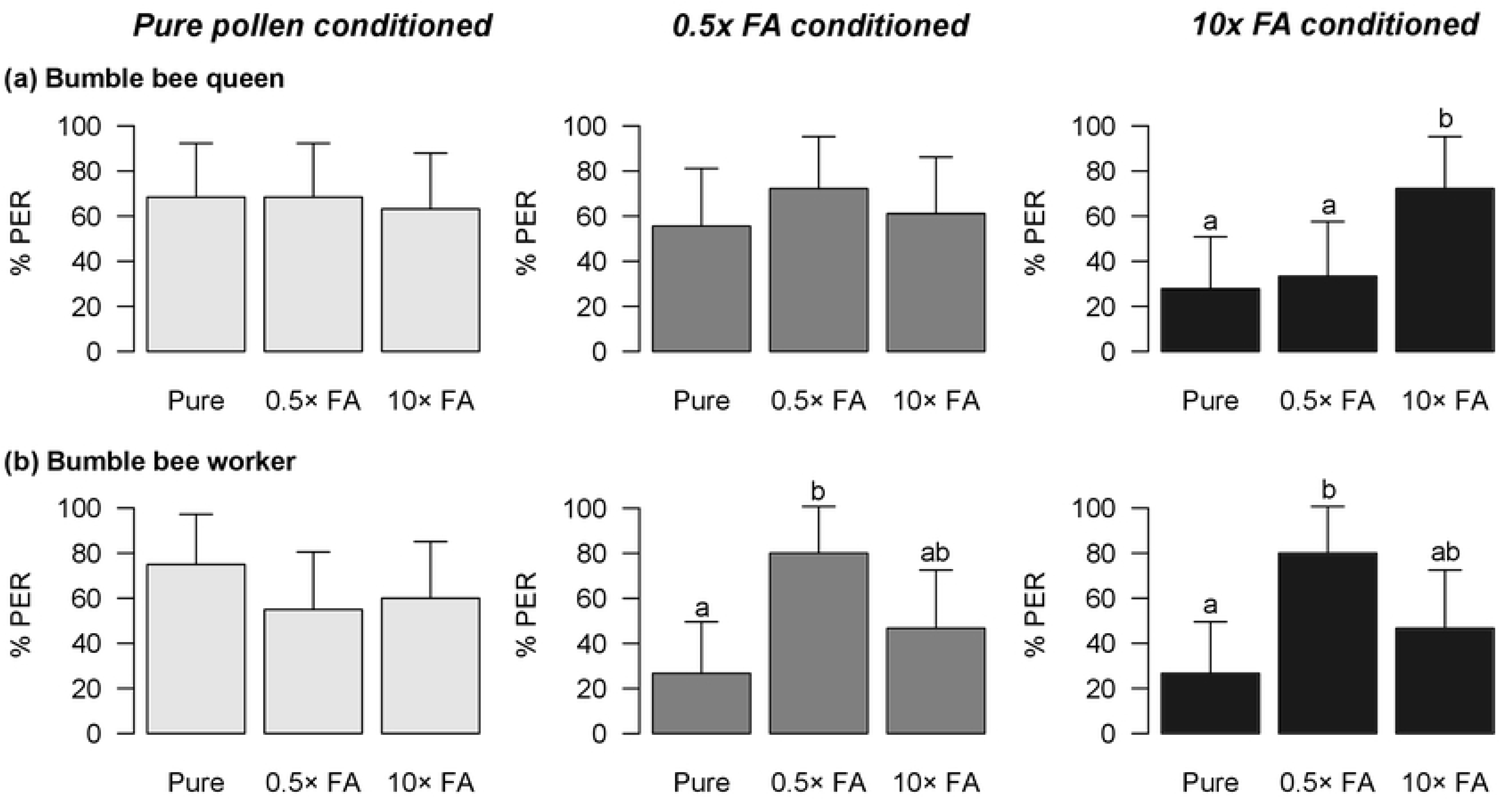
Percentage of proboscis extension responses (PER) of (a) Bumble bee queens and (b) bumble bee workers during the Test phase to the different pollen stimuli (Pure pollen with 0×FA in light gray, 0.5×FA in dark gray, and 10×FA in black). The stimulus that was used to train the different groups of bees is shown above each graph. Different letters indicate Tukey test significant differences (*P*<0.05).

**Figure 4.**
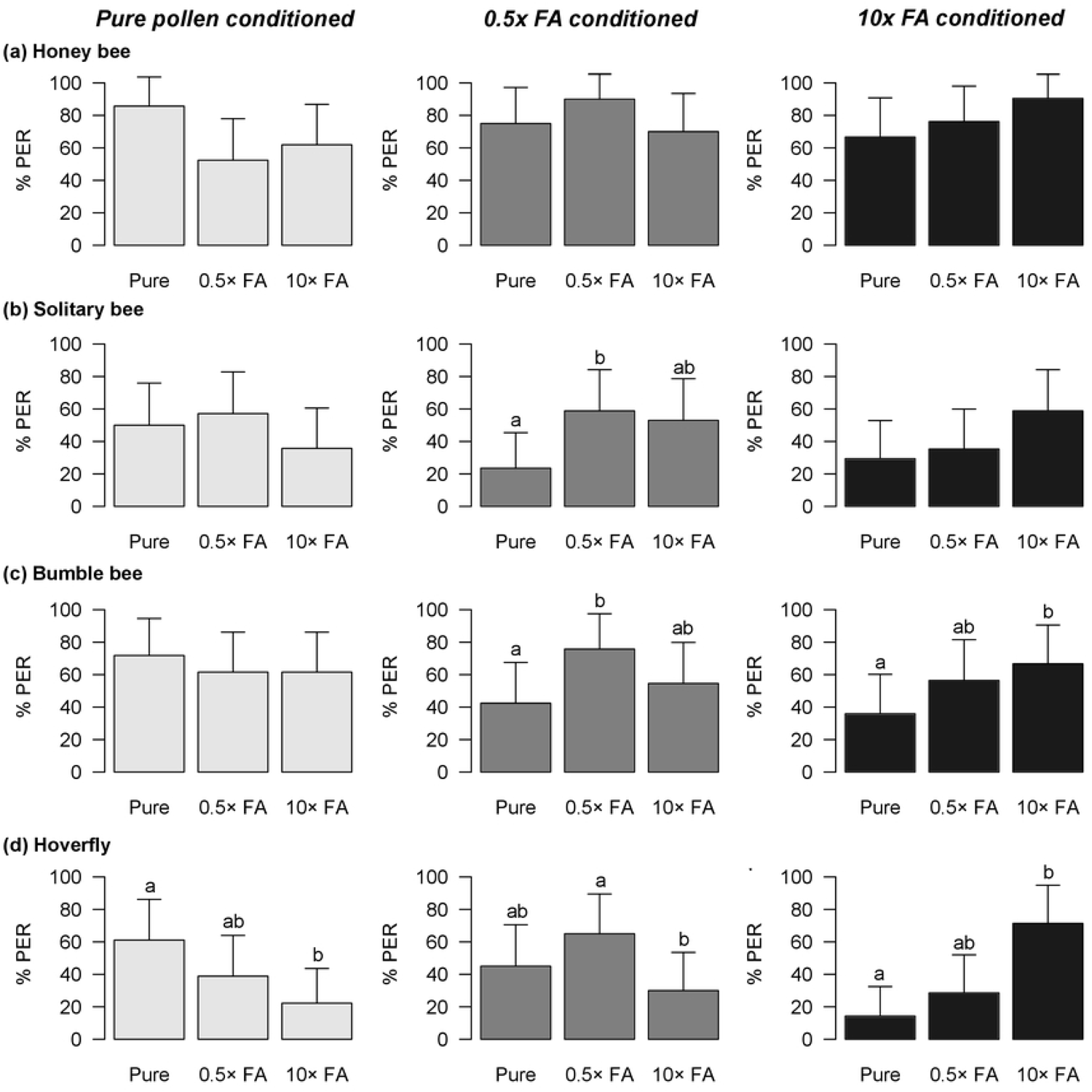
Percentage of proboscis extension responses (PER) of the four studied species for (a) honey bee, (b) solitary bee, (c) bumble bee and (d) hoverfly, during the Test phase to the different pollen stimuli (Pure pollen with 0×FA in light gray, 0.5×FA in dark gray, and 10×FA in black). The stimulus that was used to train the different groups of insects is shown at the top under (A). Different letters indicate Tukey test significant differences (*P*<0.05).

As expected, when evaluating the responses of the different pollinator species in the Test phase, we found that responses to the CS were almost always the highest (**Figure 4**). The only exception occurred in solitary bees conditioned with pure pollen. When analyzing the patterns of generalization (**Table 2**), we found in honey bees no significant differences among responses to the different stimuli, suggesting strong generalization in this species (**Figure 4A**). Solitary bees showed a similar generalization behavior as honey bees, responding similarly to the different stimuli. However, when conditioned with 0.5×FA, solitary bees generalized to the stimulus containing a higher concentration of FA but not to the pure pollen stimulus (**Figure 4B**). Bumble bees tended to generalize between the FA enriched pollen stimuli and when conditioned with enriched pollen, they responded significantly less to pure pollen (**Figure 4C**). Interestingly, when conditioned with pure pollen, bumble bees generalized fully to the other stimuli. These observations suggest that when pollen was enriched with FA it tended to overshadow the odor of the pollen compounds.

**Table 2.** Summary of the GLMM models performed to evaluate the effects of studied insects (Honey bee vs. Bumble bee vs. Hoverfly vs. Solitary bee) and pollen quality categories (Pure pollen with 0×FA vs. 0.5×FA vs. 10×FA) on the percentage of proboscis extension responses (PER) in bumble bees during Test phase. Bold lines indicate significant differences (*P*<0.05).

| Parameter | Estimate | Std. Error | z value | P value |
| --- | --- | --- | --- | --- |
| <u>Pure pollen + 0×FA conditioned</u> |  |  |  |  |
| **Intercept** | **1.23603** | **0.33869** | **3.649** | **< 0.001** |
| **Pollen quality 0.5×FA** | **-0.68537** | **0.31659** | **-2.165** | **0.030399** |
| **Pollen quality 10×FA** | **-0.86861** | **0.31630** | **-2.746** | **0.006030** |
| **Hoverfly** | **-1.10340** | **0.39175** | **-2.817** | **0.004854** |
| Solitary bee | -0.81416 | 0.41545 | -1.960 | 0.050026 |
| Bumble bee | -0.07819 | 0.33485 | -0.234 | 0.815364 |
| <u>Pure pollen + 0.5×FA conditioned</u> |  |  |  |  |
| **Intercept** | **0.8757** | **0.3568** | **2.454** | **0.014127** |
| **Pollen quality 0.5×FA** | **1.2232** | **0.3305** | **3.701** | **< 0.001** |
| Pollen quality 10×FA | 0.2406 | 0.3106 | 0.774 | 0.438662 |
| **Hoverfly** | **-1.4991** | **0.4185** | **-3.582** | **< 0.001** |
| **Solitary bee** | **-1.5668** | **0.4341** | **-3.609** | **< 0.001** |
| **Bumble bee** | **-1.0302** | **0.3827** | **-2.692** | **0.007109** |
| <u>Pure pollen + 10×FA conditioned</u> |  |  |  |  |
| Intercept | 0.6788 | 0.3901 | 1.740 | 0.081836 |
| **Pollen quality 0.5×FA** | **0.7079** | **0.3354** | **2.110** | **0.034835** |
| **Pollen quality 10×FA** | **1.6711** | **0.3637** | **4.594** | **< 0.001** |
| **Hoverfly** | **-1.8222** | **0.5224** | **-3.488** | **< 0.001** |
| **Solitary bee** | **-1.8648** | **0.5035** | **-3.704** | **< 0.001** |
| **Bumble bee** | **-1.2532** | **0.4226** | **-2.965** | **< 0.001** |

Of all the species studied, hoverflies showed the most important response pattern in the Test phase, which was quite different from that of bees. Hoverflies were the only species to always show significantly different responses among the different stimuli. In particular, it is the only species that did not generalize to both stimuli containing FA when it was conditioned with pure pollen odour. In addition, although hoverflies conditioned with the higher concentration of FA (10x) generalized to the lower concentration (0.5x FA) the reverse was not true (**Figure 4D**).

Finally, we found that for pure pollen conditioning the hoverflies’ pattern was significantly different from those of honey bees and bumble bees, by comparing the generalization behavior among the species (**Table 3**). Moreover, when insects were conditioned with enriched pollen, the generalization patterns were significantly different between honey bees and the other three species (**Table 3**).

**Table 3.** Tukey test values comparing the percentage of proboscis extension responses (PER) for different studied insects (Honey bee vs. Bumble bee vs. Hoverfly vs. Solitary bee) from the GLMMs testing the effects of studied insects and pollen quality categories on PER during Test phase. Bold lines indicate significant differences (P<0.05).

| Parameter | Estimate | Std. Error | z value | P value |
| --- | --- | --- | --- | --- |
| <u>Pure pollen + 0×FA conditioned</u> |  |  |  |  |
| **Hoverfly vs. Honey bee** | **-1.10340** | **0.39175** | **-2.817** | **0.0245** |
| Solitary bee vs. Honey bee | -0.81416 | 0.41545 | -1.960 | 0.2010 |
| Bumble bee vs. Honey bee | -0.07819 | 0.33485 | -0.234 | 0.9955 |
| Solitary bee vs. Hoverfly | 0.28924 | 0.42221 | 0.685 | 0.9017 |
| **Bumble bee vs. Hoverfly** | **1.02521** | **0.34436** | **2.977** | **0.0153** |
| Bumble bee vs. Solitary bee | 0.73597 | 0.37112 | 1.983 | 0.1920 |
| <u>Pure pollen + 0.5×FA conditioned</u> |  |  |  |  |
| **Hoverfly vs. Honey bee** | **-1.49914** | **0.41851** | **-3.582** | **0.00201** |
| **Solitary bee vs. Honey bee** | **-1.56683** | **0.43413** | **-3.609** | **0.00187** |
| **Bumble bee vs. Honey bee** | **-1.03020** | **0.38273** | **-2.692** | **0.03522** |
| Solitary bee vs. Hoverfly | -0.06769 | 0.39572 | -0.171 | 0.99820 |
| Bumble bee vs. Hoverfly | 0.46894 | 0.34057 | 1.377 | 0.51100 |
| Bumble bee vs. Solitary bee | 0.53663 | 0.35944 | 1.493 | 0.43882 |
| <u>Pure pollen + 10×FA conditioned</u> |  |  |  |  |
| **Hoverfly vs. Honey bee** | **-1.82218** | **0.52241** | **-3.488** | **0.00274** |
| **Solitary bee vs. Honey bee** | **-1.86480** | **0.50346** | **-3.704** | **0.00141** |
| **Bumble bee vs. Honey bee** | **-1.25317** | **0.42261** | **-2.965** | **0.01541** |
| Solitary bee vs. Hoverfly | -0.04262 | 0.51904 | -0.082 | 0.99980 |
| Bumble bee vs. Hoverfly | 0.56901 | 0.42733 | 1.332 | 0.53760 |
| Bumble bee vs. Solitary bee | 0.61163 | 0.42403 | 1.442 | 0.46756 |

## Discussion

To our knowledge, this is the first comparative study of learning behavior (associative learning) between multiple insect pollinators using the FMPER procedure including a native solitary bee and a hoverfly species. We studied the perception, learning and generalization ability among pollen enriched with different concentrations of stearic acid (a fatty acid) in four species of insect pollinators. We demonstrated that all pollinator species were able to associate the different concentrations of enriched pollen with a reward, since the PER level increased over the course of the trials in the conditioning phase of the experiment. Also, our results showed that learning performances differ between castes of bumble bees (*B. terrestris*), as well as between honey bees and other species.

Bumble bee queens showed better learning performances than workers for all pollen stimuli. Similarly, Muth *et al.* (2021) [42] demonstrated that *B. vosnesenskii* queens and workers differed in their ability to learn a color-reward association and found that queens performed better than workers. Ruedenauer *et al.* (2020) [37] also showed that workers of *B. terrestris* could perceive and learn pollen enriched with fatty acids but do not prefer pollen with higher concentrations of fatty acids, which produces a decrease in their survival. The consequences of learning are not the same for the different castes of the colony. If a queen does not learn quickly (specially at the moment of start a new colony) the future development may be at risk, while in the case of workers the consequences are less severe mainly affecting the individual engaged in cooperative work. From a nutritional perspective, if the learned stimulus is not nutritionally adequate, the consequences for the colony may be worse if this occurs in queens than in workers. Perhaps for this reason, the pattern of generalization differs between castes. Therefore, it is possible that queens have better learning capabilities than workers due to the implications of their life cycle, their nutritional requirements (especially after emerging from hibernation) and for the success of the future colony. Perhaps, workers learn slower but because they have more time to learn and exploit better resources and being more specifics. In future studies it would be of great interest to evaluate the survival of queens with foods of different nutritional qualities.

When we evaluated learning success among the four pollinator species studied, we found significant differences in the species’ performances but the stimuli are learned in a similar way. In a recent study performed by Kandori *et al*. (2021) [28] the authors explored differences in the color-learning rates among eight pollinator species of three insect orders (Hymenoptera, Diptera, and Lepidoptera). They demonstrated that learning performances while foraging for flowers differed among these species and calculating learning rates, they established a gradation among them: the highest learning rate was in *Bombus ignitus*, followed by the two larger butterflies (*Idea leuconoe* and *Argyreus hyperbius*), the two smaller butterflies (*Pieris rapae* and *Lycaena phlaeas*), and the remaining species (*E. cerealis, E. tenax,* and the solitary bee *Osmia orientalis*). They argued that learning differences are possibly due to the degree of eusociality, body size and degree of nutritional specialization of the species. We found a similar gradation in which eusocial bees were first, followed by the hoverfly and the solitary bee. At the same time, this gradation of learning performances is also in agreement with the hypothesis proposed by Collado *et al*. (2021) [26], which argued that bees’ body size largely determines learning ability. We considered that, it will be necessary to compare a wider range of species to disentangle the respective effects of size and social lifestyle on learning capabilities. Concerning hoverflies, Lunau *et al*. (2018) [57] demonstrated that *E. tenax* has certain limitations when learning visual stimuli. Our results are on the same line and indicate a possible learning limitation in this species, which only showed good learning performances for the pollen with the low concentration of fatty acid. We consider that the differences that we found may also depend on the combination of foraging strategies and the adaptive value of this behavior, qualities that are specific to each species and that are strongly related to their natural history, nutritional requirements, feeding preferences and neural development [58–62].

Parallel to the learning patterns that we found, the ability to generalize also differed between the different insect species. According to our results, there is a generalization gradient between insect species, from honey bees that generalized the most, to the solitary bee, followed by bumble bees and finally the hoverfly. Honey bees differed remarkably from the other species, as they were the only ones that generalized among all three stimuli regardless of fatty acid concentration. The native solitary bee, *R. mutabilis* had a similar pattern of generalization as honey bees, and perceived almost all stimuli as similar. Until now, despite being very abundant in Chile and the northwest of Argentinean Patagonia, as well as a good pollinator of several native and exotic species*, R. mutabilis* has been studied very little [39, 63–64]. Indeed, here we present a pioneering study of its learning capability.

The three hymenoptera species perceive the pollen enriched with a high concentration of stearic acid as a salient stimulus and were able to generalize stimuli much more than the hoverfly. Hoverflies were the species that showed the most peculiar learning pattern since they learned the intermediate FA concentration (0.5×FA) better than the two other stimuli, especially the higher FA concentration (10x FA). Previous research [57] also demonstrated that *E. tenax* possess some learning limitations depending on the type of cues used during the conditioning procedure. Regarding generalization behavior, this is the only species that did not respond to all the other stimuli when they were trained with pure polen. This species thus seems to be relatively specific in its responses to olfactory stimuli, to test this question it would be interesting to perform a FMPER protocol of differential learning, in which one stimulus is explicitly rewarded (CS+) while another is explicitly unrewarded [65].

Under natural conditions, the ability to generalize two stimuli might contribute to improve foraging activities but depending on the qualities of the generalized stimuli the adaptive implications could be beneficial, neutral or detrimental. The fact that pollen highly enriched with fatty acids was generalized can have a negative impact on colony survival, especially if we consider that the consumption of large amounts of fatty acids negatively affects survival, as has been shown in other research [37]. Determining the pattern of pollen generalization will provide a better understanding of the ecological implications of this behavior; since this behavior play a key role in the immediate and future success of the individual and colony, and consequently in the services and products they provide [66]. For this reason, it is also necessary to carry out complementary studies on the ability to discriminate between similar stimuli (for example, other fatty acids) at different concentrations, as well as include survival and reproductive experiments.

Finally, we would like to point out that despite the fact that this type of experimental protocol performed in the field on freely moving individuals (FMPER) implies a greater work effort than traditional Lab approaches with harnessed individuals, we consider that they are of great importance since they allow to tackle the behavior of native insects. As they are less detrimental to the welfare of the experimental animals, the study subjects can be rapidly released after the experiment. Studying the foraging behavior, learning capabilities and the adaptive value of learning in wild pollinator insects provides valuable information for the conservation of these species and also for beekeepers and agricultural producers allowing them to develop nutritional strategies appropriate to the needs of their agro-ecosystems.

## Acknowledgements

We are grateful to Dra. Paula Crego (Universidad Nacional del Comahue -CRUB) and to Ing. Guillermo Huerta (INTA EEA Bariloche) for their helpful discussions on this project. We also thank Joshua Taylor for his valuable comments in previous versions of the manuscript. Finally, Dr. Ana Laura would like to especially thanks to her mother Norma Telechea for her support during the sampling of insects carried out in times of confinement due to the COVID-19 pandemic.

## Funding

This work was financed by Fondo para la Investigación Científica y Tecnológica-FONCyT-Argentina (PICT 2018-0952) and by Instituto Nacional de Tecnología Agropecuaria-INTA-Argentina (PE 074).

## Disclosure

The authors declare that the research was conducted in the absence of any commercial or financial relationships that could be construed as a potential conflict of interest.

## Data accessibility

Should the manuscript be accepted, the data that support the findings of this study will be openly available through the figshare repository https://doi.org/xxxx/m9.figshare.xxxxx (Pietrantuono et al., xxx).

## Author Contribution

**Conceptualization**: Ana Laura Pietrantuono, Fabrice Requier

**Data curation**: Ana Laura Pietrantuono

**Formal analysis**: Fabrice Requier

**Funding acquisition**: Ana Laura Pietrantuono

**Investigation**: Ana Laura Pietrantuono, Valeria Fernández-Arhex

**Methodology**: Ana Laura Pietrantuono, Fabrice Requier

**Project administration:** Fabrice Requier

**Supervision**: Fabrice Requier, Jean-Christophe Sandoz

**Visualization**: Fabrice Requier

**Writing – original draft**: Ana Laura Pietrantuono, Fabrice Requier

**Writing – review & editing**: Ana Laura Pietrantuono, Valeria Fernández-Arhex, Jean-Christophe Sandoz, Fabrice Requier

## Notes

### Competing Interest Statement

The authors have declared no competing interest.

